# Integrated kidney and urine proteomics define encrypted antimicrobial peptides as effectors of host defence in human pyelonephritis

**DOI:** 10.64898/2026.04.10.717476

**Authors:** Lars Borgards, Hannah Voss, Stephanie Tautges, Bente Siebels, Ekaterina Pylaeva, Philippa Spangenberg, Devon Siemes, Christoph Krisp, Jessica Schmitz, Lisa Cinkul, Damilola Akinyemi, Annika Hilger, Paulina Szymczak, Anne Aust, Jan H Bräsen, Ewa Szczurek, Cesar de la Fuente-Nunez, Jadwiga Jablonska, Oliver Soehnlein, Ulrich Dobrindt, Hartmut Schlüter, Sibylle von Vietinghoff, Florian Wagenlehner, Daniel R. Engel, Olga Shevchuk

**Affiliations:** Department of Immunodynamics, Institute for Experimental Immunology and Imaging, University Hospital Essen, Essen, Germany; Section Mass Spectrometry and Proteomics, University Medical Center Hamburg-Eppendorf (UKE), Hamburg, Germany; Department of Otorhinolaryngology, Head and Neck Surgery, University Hospital Essen, University of Duisburg-Essen, Essen, Germany; German Cancer Consortium (DKTK) partner site Düsseldorf/Essen; Essen, Germany; Nephropathology Unit, Institute for Pathology, Hannover Medical School, Hannover; Germany; Clinic for Urology, Pediatric Urology and Andrology, Justus-Liebig-University Giessen, Giessen, Germany; Institute of Experimental Pathology (ExPat), Center for Molecular Biology of Inflammation (ZMBE), University Hospital Münster, University of Münster, Münster, Germany; Institute of Hygiene, University of Münster, Münster, Germany; Institute of AI for Health, Helmholtz Zentrum München, German Research Center for Environmental Health, Neuherberg, Germany; Institute of Informatics, Faculty of Mathematics, Informatics and Mechanics, University of Warsaw, Warsaw, Poland; Machine Biology Group, Departments of Psychiatry and Microbiology, Institute for Biomedical Informatics, Institute for Translational Medicine and Therapeutics, Perelman School of Medicine, University of Pennsylvania, Philadelphia, Pennsylvania, United States of America; Departments of Bioengineering and Chemical and Biomolecular Engineering, School of Engineering and Applied Science, University of Pennsylvania, Philadelphia, Pennsylvania, United States of America; Department of Chemistry, School of Arts and Sciences, University of Pennsylvania, Philadelphia, Pennsylvania, United States of America; Penn Institute for Computational Science, University of Pennsylvania, Philadelphia, Pennsylvania, United States of America; Nephrology Section, Medical Clinic 1, University Hospital Bonn, Rheinische Friedrich-Wilhelms University, Bonn, Germany

**Keywords:** pyelonephritis, antimicrobial peptides, urinary and renal proteome, urinary peptidome, human kidney, encrypted AMPs, innate immunity

## Abstract

Antimicrobial peptides (AMPs) are key effectors of host defence, however, their functional deployment across renal tissue and urine in pyelonephritis (PN) remains incompletely understood. Here, we integrate kidney and urine proteomics with urinary peptidomics and computational prediction to define AMP organisation and function. Proteomic analysis indicated coordinated induction of multiple AMPs in infected kidneys. These patterns were recapitulated in the urinary proteome, where AMP abundance correlated with leukocyte counts. Multiplex immunofluorescence microscopy localised these AMPs to myeloid cells, identifying them as central effector sources. Importantly, analysis of the urine peptidome revealed multiple encrypted AMPs (EPs), which arise from proteolytic processing of precursor proteins. To systematically assess their relevance for host defence, we applied an ensemble of machine learning-based predictors to prioritise candidates with activity in the urinary environment. This approach identified several potential EPs, among which the S100A12-derived peptide Calcitermin was confirmed in patient urine. Furthermore, it exerts antibacterial activity against uropathogenic *E. coli* (UPEC) and modulates myeloid cell responses. Together, these findings define a coordinated and compartmentalised AMP defence programme in human PN that extends beyond increased peptide expression, highlighting EPs as functionally relevant effectors with therapeutic potential.

## Introduction

Pyelonephritis (PN) is a common upper urinary tract infection characterized by inflammation of the renal parenchyma and is associated with substantial morbidity. Potential complications of PN include hypertension, proteinuria, renal fibrosis, kidney failure, and urosepsis^1,2^. Although, PN is currently treated with antibiotics, multidrug-resistant uropathogens represent an increasing clinical challenge^3^, with 17 – 36% of urosepsis being caused by pathogens resistant against the initial empiric antibiotic treatment^4^. Therefore, optimized treatment strategies and the development of novel therapeutic approaches are required.

The innate immune response plays a central role in protecting the urothelium and renal parenchyma from pathogens^5^. Both leukocytes and epithelial cells mediate innate immunity through direct phagocytosis of the pathogen and the release of soluble factors, including components of the complement system, chemokines, cytokines, antimicrobial peptides (AMPs)^6–8^, and recently described encrypted AMPs (EPs) – short antimicrobial fragments derived from larger precursor proteins^9^. In addition to the direct inactivation of bacteria, AMPs modulate the immune response, enhance chemokine production, regulate epithelial cell differentiation, and alter angiogenesis^6^. Thus, AMPs have gained increased attention as potential prognostic markers and candidates for novel therapeutics^10^. Several AMPs have been defined in the context of urinary tract immunity, including cathelicidin antimicrobial peptide (LL-37 or CAMP)^11^, various α- and β-defensins^11–14^, RNase 7^15^, pentraxin-related protein 3 (PTX3)^16^, uromodulin (UMOD)^17,18^, and metal-binding AMPs, such as lipocalin-2 (LCN2)^18–20^, lactotransferrin (LTF)^21^, and the S100A8/S100A9 heterodimer calprotectin^22–25^. The epithelial cells of the urinary tract were defined as primary sources of AMPs, complementing contributions from innate immune cells. Furthermore, α-intercalated cells of the collecting duct and thick ascending limb cells play a key role in the production and release of antimicrobial factors in the kidney^8,26^.

Given the central role of immune and antimicrobial responses are central to PN, kidney proteomics may identify disease-specific, targetable biomarkers and therapeutic candidates. In this context, profiling the urinary proteome in PN patients is of particular interest, as urine represents a non-invasive diagnostic tool that reflects the secreted proteome of the tubular fluid.

This, in turn, recapitulates the local defensive proteome of the renal tubular system and surrounding interstitial tissue—the primary sites of infection—thereby providing direct insights into host–pathogen interactions within the kidney. Although the function of individual AMPs has been delineated in the context of UTI, the global AMP landscape in PN remains unexplored. While data mining—computational analysis of large datasets—using predictive discriminators has shown promising initial results in identifying EPs based on amino acid sequence analyses of the human proteome, these peptides are rarely verified for their presence and functional relevance *in vivo*, particularly under pathological conditions. Building on this concept, we recently established a strategy for global *in vivo* profiling of antimicrobial proteins and EPs in UTI^27^. In the current study, we combine proteomic and peptidomic analyses of renal tissue and urine from PN patients to interrogate the organisation and functional deployment of AMP defences during infection. These analyses indicate strong induction of immune cell–derived AMPs in infected kidneys and their accumulation in urine. To systematically assess antimicrobial relevance across the detected peptides, we applied an ensemble of machine learning classifiers and regressors, including both broad-spectrum AMP predictors and strain-specific models, prioritising peptide candidates with predicted activity against uropathogenic bacteria. Among these, we focused on Calcitermin as an EP candidate predicted to exert antimicrobial activity in the urinary environment. Its presence in patient urine was confirmed by targeted proteomics, and its antibacterial activity against uropathogenic *E. coli*, as well as its immunomodulation of the myeloid compartment, was validated *in vitro*. Together, these integrative approaches define a disease-specific framework of AMP regulation in human PN and highlight bioactive EP effectors with potential therapeutic relevance.

## Materials and methods

### Kidney samples of PN patients

PN nephrectomy specimens (*n*=10; 20% men; mean age: 51.8 years; range: 22 – 76 years) and healthy regions from tumor resections (*n*=10; 30% men; mean age: 48.4 years; range: 7 – 82 years) were used from the archives of the Nephropathology Unit at Hannover Medical School (MHH). Ethic MHH: No. 10183_BO-K_2022. The underlying diagnoses in patients undergoing nephrectomy are summarized in Supplementary Table 1.

### Urine samples of PN patients

Urine was collected from acute PN patients (*n*=30) and healthy control donors (*n*=10) by the Clinic for Urology, Pediatric Urology and Andrology of the Justus-Liebig-University Giessen. The ethical approval has been received by the independent ethics commission of the University Giessen, Germany (Ethical vote 280/20) on January 6^th^, 2021. The clinical data of PN patients and control urine donors are summarized in Supplementary Table 3. The cases of acute PN were clinically diagnosed, following the FDA’s definitions used for studies in the indication “Developing drugs for treatment. Guidance for industry, June 2018” (https://www.fda.gov/media/71313/download). Patients admitted to the urological department at the University Hospital Giessen with at least two of the following signs or symptoms were included: chills, rigors, or warmth associated with fever; flank pain; nausea or vomiting; dysuria; urinary frequency; urinary urgency; costo-vertebral angle tenderness upon physical examination; or a urine specimen with evidence of pyuria. Risk factors for recurrent UTI were categorized by urologists according to the ORENUC classification system^28^. Urine samples were centrifuged at 5,000 × g for 10 min at 4 °C, and the supernatant was filtered through a 0.22 µm sterile filter (Merck Millipore, Burlington, USA).

### Enzyme-linked immunosorbent assay (ELISA)

Urinary S100A8/S100A9 (Immundiagnostik, K 6928), S100A12 (Thermo Fisher, EH170RB), cathepsin G (antibodies-online, ABIN6954455), azurocidin-1 (antibodies-online, ABIN4881912), and UMOD (BioVendor, RD191163200R) were quantified by ELISA according to the manufacturers’ protocols. Absorbance was measured at 450 nm (iMark™, Bio-Rad), and values were normalized to creatinine.

### Patient sample processing for LC–MS/MS proteome and peptidome

For renal proteomics, FFPE kidney tissue sections (60 µm) were prepared as previously described^29^. For urine proteomics, soluble proteins were extracted using acetone precipitation according to Pichler et al.^30^.

For native peptidomics, the soluble urine fraction was first filtered to retain molecules <50 kDa using Pierce™ Protein Concentrators PES, 50K MWCO (0.5–100 mL; Thermo Fisher Scientific, Waltham, USA), according to the manufacturer’s instructions. The flow-through was subsequently filtered to retain molecules <10 kDa using Pierce™ Protein Concentrators PES, 10K MWCO (0.5–100 mL; Thermo Fisher Scientific, Waltham, USA). Molecules <10 kDa were dried in a SpeedVac concentrator.

Both native and tryptic peptides were desalted using Pierce™ C18 Spin Tips & Columns (Thermo Fisher Scientific, Waltham, USA) according to the manufacturer’s instructions, dissolved in 40 µL of 0.1% formic acid (FA). Peptide concentrations were estimated using a NanoDrop Microvolume Spectrometer (Thermo Fisher Scientific, Waltham, USA) at 205 nm.

For absolute quantification, synthetic light (VAIALKAAHYTHKE) and heavy-labeled ((VAIALKAAHYTHK (Lys 13C6; Lys 15N2) E) Calcitermin were obtained from JPT Peptide Technologies (Berlin, Germany) at ≥95% purity. Calibration standards were prepared by dissolving 3.125–200 fmol of light Calcitermin in 15 ng/µL of pooled native urinary peptides from PN patients and controls. All samples and calibration standards were spiked with 25 fmol heavy-labeled Calcitermin for normalization. For spectral library generation, 200 fmol each of light and heavy Calcitermin were dissolved in 0.1% FA.

### LC-MS/MS data acquisition

For kidney tissue samples, LC–MS/MS measurements were performed on Orbitrap Fusion (Thermo Fisher Scientific) coupled to a nano-UPLC system (Dionex Ultimate 3000, Thermo Fisher Scientific), as previously described^29^.

For urinary proteome and peptidome analysis, 300 ng of tryptic or native peptides were loaded onto an Evotip (Evosep Biosystems, Odense, Denmark) according to the manufacturer’s instructions. Peptide separation was carried out on an Evosep One system (Evosep Biosystems) using the predefined “30SPD” method, on a PepSep Series C18 column (15 cm × 150 µm inner diameter, 1.5 µm particle size; Bruker Daltonics, Billerica, MA, USA). Eluting peptides were introduced into a timsTOF HT mass spectrometer (Bruker Daltonics) via a CaptiveSpray source at 1,600 V.

Urinary proteomics was performed using data-independent acquisition (DIA) parallel accumulation-serial fragmentation (PASEF) on a timsTOF mass spectrometer as described previously^31^.

Urinary native peptidomics was performed using data-dependent acquisition (DDA) - PASEF mode, with ten PASEF scans per topN acquisition cycle and a total cycle time of 1.17s. The accumulation and ramp time for the dual TIMS analyzer were set to 100 ms at a ramp rate of 9.42 Hz. Scan ranges were set to *m*/*z* 100\-1,700 for the mass range and 1 / *k*0 0.60-1.60 V s^−1^ cm^−2^ for the mobility range at both MS levels. Suitable precursor ions for PASEF-MS/MS were selected based on their positions in the *m*/*z*-ion mobility plane in real time from TIMS-MS survey scans. Precursors that reached a ‘target value’ of 20,000 AU were dynamically excluded for 0.4 min. The width of quadrupole isolation was set to 2 Th for *m*/*z* < 700 and 3 Th for *m*/*z* > 800. The CE was ramped linearly as a function of the mobility, from 52 eV at 1.6 V s^−1^ cm^−2^ to 20 eV at 0.6 V s^−1^ cm^−2^.

For absolute quantification, the mass spectrometer was operated in parallel reaction monitoring (PRM)-PASEF mode. For MS1 scanning, similar settings as described for DDA-PASEF measurements were selected. MS2 was only triggered between 5.5 and 7.5 minutes for precursors within a mass to charge range of 562 to 850 m/z and an ion mobility of 0.82 to 1-16 1/KO, corresponding with the double and triple charged versions of heavy and light Calictermin.

### LC-MS/MS data analysis for kidney tissue

LC-MS/MS data were searched using the Sequest algorithm in Proteome Discoverer (v2.41.15, Thermo Fisher Scientific) against a reviewed human Swissprot database (April 2021; 20,365 entries). Carbamidomethylation of cysteine was set as a fixed modification, while methionine oxidation, N-terminal glutamine pyroglutamate formation, and protein N-terminal acetylation were included as variable modifications. Up to two missing tryptic cleavages were allowed, and peptides of 6-144 amino acids were considered. FDR < 0.01 was set for peptide and protein identification. Quantification was performed using the Minora Algorithm, followed by log_2_-transformation of protein abundancy and LOESS normalization^32^.

For statistical analysis, 2,562 proteins detected in at least one condition with ≥2 biological replicates were included. For each protein, fold change, p-value, and signal-to-noise ratio were calculated, and p-values were adjusted for FDR using the Benjamini–Hochberg method (q-value). Proteins with |log2(FC)| > 2 and q < 0.05 were considered significantly regulated; 160 (6.2%) were upregulated and 507 (19.8%) downregulated. Cytoscape (Cytoscape_v3.8.0, ClueGO_v2.5.7; gene ontology databases from 08.05.2010; min GO level = 3; max GO level = 8; number of genes = 3; min percentage = 3.0; kappa score threshold = 0.4) was used for functional enrichment analysis.

The Pearson correlation Clusterheatmap was generated in Python with plotly (v5.5.0) using 1273 proteins with a *q*-value < 0.05. Pearson correlation was calculated in pandas (v1.3.5), and the clustering was computed with scipy (v1.7.3). Regression plots were generated using the regplot method of the Seaborn (v.0.11.2) data visualization library for Python.

### LC-MS/MS data analysis for urinary proteome and peptidome

For proteomics, LC-MS data were processed using the directDIA workflow in Spectronaut (v19.1.240806.626, Biognosys), applying BSG factory settings (Trypsin/P; fixed modification: carbamidomethyl (C); variable modifications: acetyl (protein N-term) and oxidation(M)), against the reviewed human UniProt/Swissprot FASTA database (March 13, 2024; 20,418 entries). Quantification was performed at the MS2 level with imputation and cross-run normalization disabled. Protein abundances were exported to R (v 4.1.2), log_2_-transformed, and normalized by subtracting the mean abundance per sample for each protein quantified. Missing values were imputed under a missing-not-at-random assumption using a protein-wise normal distribution (downshift 1.3, width 0.3). Differential abundance was assessed by Student’s t-test with significance defined as FDR (Benjamini Hochberg) < 0.05 and ≥1.5-fold change. Principal component analysis was performed on normalized and imputed data.

For native peptidomics, LC-MS raw data was processed in MaxQuant (v2.6.7) using the Andromeda search engine with unspecific digestion against the same UniProt/Swissprot FASTA database. Protein N-terminal acetylation and methionine oxidation were set as variable modifications. Peptides longer than six amino acids and below 10 kDa were considered, with match-between-runs enabled.

For absolute quantification, a spectral library was generated from 200 fmol heavy and light Calcitermin in 0.1% FA. The raw data were processed with FragPipe (v23.1) using MSFragger and SpecLib and searched against a custom Calcitermin FASTA database supplemented with common contaminants under the assumption of unspecific proteolytic digestion^33,34^. From the resulting library, fragmentation patterns corresponding to triply charged light (m/z 563.317) and heavy (m/z 566.317) Calcitermin ions were extracted and subsequently imported into Skyline (v25.1, University of Washington, Seattle, USA)^35^.

For absolute quantification using Skyline, the following transition settings were applied: Triply charged precursor ion; charges of 1 or 2 for product ions; y, b, and precursor ions as considered ion types. From each precursor ion, the top 10 product ions were considered with an ion match tolerance of 0.5 m/z. At the instrument level, a method match tolerance of 0.05 m/z was selected. TOF was chosen as a mass analyzer at a resolution of 30.000 at MS1 and MS2 levels, respectively. Only scans within 5 minutes of MS/MS IDs were considered. For quantification, a linear regression fit was used for the calibration curve. Calcitermin intensities were normalized to the intensities of the heavy-labeled Calcitermin. The LOD was estimated by blank intensities + 2 * SD. For LOQ, a Coefficient of Variation (CV) of a maximum 20 % was accepted. Calcitermin intensities for patient samples were furthermore normalized to the measured peptide amount.

### Antimicrobial activity prediction and scoring

Peptide sequences were analyzed for antimicrobial potential using published machine learning classifiers and regressors, grouped by prediction type. General AMP identification was performed using six classifiers: AMPeppy^36^, AMPlify^37^, AMPScanner^38^, HydrAMP-AMP-classifier^39^, OmegAMP broad classifier^40^, and SenseXAMP-classifier^41^.

Strain- and species-specific activity was predicted using the HydrAMP-MIC-classifier for *E. coli* and 13 OmegAMP models trained on experimentally validated datasets for defined bacterial strains and species. OmegAMP represents a conservative prediction system with a low FDR, but limited sensitivity compared to other methods.

Predicted minimum inhibitory concentrations (MICs) were obtained using Deep-AMP models (for Gram-positive and Gram-negative bacteria) and SenseXAMP regressors (for *Staphylococcus aureus* and *E. coli*). For downstream analysis, predictions were summarized as: (i) classifier rank, representing the number of broad AMP classifiers predicting a peptide as antimicrobial (maximum rank = 5); (ii) OmegAMP rank, representing the number of strain- and species-specific OmegAMP classifiers predicting a peptide as active (maximum rank = 13); (iii) HydrAMP-MIC and OmegAMP broad-spectrum predictions for discriminating active from inactive peptides; and (iv) predicted MIC values from SenseXAMP and Deep-AMP models.

### Immunofluorescence microscopy and image analysis

Deparaffinization, antigen retrieval, and staining of 8 µm FFPE human kidney tissue sections on Superfrost® glass slides were performed according to the “MACSima™ Imaging Cyclic Staining (MICS) Sample preparation protocol for FFPE tissue” (v1.1, Jun 2021) from Miltenyi Biotec. Cyclic immunofluorescence imaging was performed as described previously^42^. Antibodies targeted AQP1 (Sigma-Aldrich, HPA019206), NKCC2 (Atlas Antibodies, HPA014967), CD14 (Miltenyi Biotec, 130-110-576), CD66b (Miltenyi Biotec, 130-122-922), AZU1 (Atlas Antibodies, HPA075964), CTSG (Atlas Antibodies, HPA047737), and S100A8 (Sigma-Adrich, HPA024372). Afterwards, the slide was photobleached, washed three times in PBS for 3 min each, and stained for S100A12 (Atlas Antibodies, HPA002881). Corresponding images were acquired on a Zeiss AxioScan.Z1.

Acquired images were stitched and preprocessed either using the MACS iQ View Analysis (Miltenyi Biotec) or ZEN Blue (Carl Zeiss GmbH) software. For the co-registration of images acquired on the MACSima system and Zeiss AxioScan.Z1, the initial transformation was performed in Fiji (rotation and cropping). For precise registration, symmetric normalization (SyNRA) with rigid, affine, and deformable transformation, and Mattes mutual information as the optimization metric implemented in the ANTs library was used^43^. The microscopy image was downsampled, using linear interpolation, to the same spatial dimensions using the Python package ImgAug (https://github.com/aleju/imgaug). Downstream image analysis was conducted in QuPath (v0.5.1). Neutrophils and monocytes were identified using CD66b^+^ and CD14^+^ CD66b^-^ staining, respectively, and analyzed for co-localization with AMPs. Additionally, AQP1^+^ and NKCC2^+^ signals were annotated as proximal and distal tubules, respectively, and evaluated for co-localization.

### Antimicrobial assays

M9 minimal medium^44^ supplemented with 0.4 % D-glucose and 0.4 % casamino acids was adjusted to pH 5.0 or 7.0. Single colonies of UPEC strain 536 were inoculated in 2 m L M9 medium and incubated at 37 °C and 180 rpm overnight. The optical density (OD_595nm_) was adjusted to 0.01 in a final volume of 150 µL of M9 medium supplemented with titrated concentrations of Calcitermin in a 96-well F-bottom microplate. Over the next 24 h, the optical density was monitored with a TECAN Infinite F200 microtiter plate reader as previously described^45^. After 24 h, samples with a final OD_595nm_ above 0.8 were diluted 1:4 in M9 medium for endpoint measurement. To assess bacterial viability, selected samples were serially diluted in 0.9% NaCl and plated on LB agar to determine colony-forming units (CFUs).

### Preparation and functional analysis of primary human monocytes and neutrophils

Peripheral blood was collected into 3.8% sodium citrate anticoagulant monovettes (Sarstedt, cat. no. 02.1067.001) and diluted 1:1 with PBS (Gibco, cat. no. 14040133). Samples were subjected to density gradient centrifugation using Biocoll (density 1.077 g/mL; Merck, cat. no. L6115) for 30 min at 300 × g (without brake) at 20°C. The mononuclear cell fraction was collected, resuspended in buffer (PBS supplemented with 0.5% bovine serum albumin (R&D Systems, cat. no. 5217) and 2 mM EDTA (Sigma-Aldrich, cat. no. E9884)), and centrifuged at 300 × g at 4°C. This washing step was repeated three times to minimize platelet contamination. CD14+ monocytes were isolated from the mononuclear fraction using immunomagnetic separation (CD14 MicroBeads, human; Miltenyi, cat. no. 130-097-052) according to the manufacturer’s instructions. Isolated monocytes (purity ~95%) were maintained on ice in RPMIc medium (RPMI 1640 (Gibco, cat. no. 11875093) supplemented with 10% fetal bovine serum (PAN Biotech, cat. no. P30-3031) and 1% penicillin–streptomycin (Gibco, cat. no. 15140122)) for up to 1 h.

Neutrophils were isolated from the polymorphonuclear fraction by sedimentation over 1% polyvinyl alcohol (Sigma-Aldrich, cat. no. 360627) for 30 min at 20°C, followed by hypotonic lysis of erythrocytes using 0.2% NaCl for 1 min at 20°C and restoration of isotonicity with 1.2% NaCl. Isolated neutrophils (purity ~95%) were resuspended in RPMIc medium at 20°C at a final concentration of 4 × 10^6^ cells/mL. Cells were seeded at 1 × 10^6^ cells/mL in RPMIc medium and incubated with 0, 4, or 16 µM peptide for 18 h at 37°C in 5% CO_2_.

Spontaneous apoptosis was assessed using an Annexin V/7-aminoactinomycin D (7-AAD) apoptosis detection kit (BD, cat. no. 559763). The proportions of viable (Annexin V^−^/7-AAD^−^), early apoptotic (Annexin V^+^/7-AAD^−^), and late apoptotic/dead (Annexin V^+^/7-AAD^+^) cells were determined within the single-cell population.

Phagocytosis was evaluated using the pHrodo™ 647 E. coli BioParticles Phagocytosis Kit (Thermo Fisher Scientific) according to the manufacturer’s instructions. Cells were incubated at 37°C with 5% CO_2_ for 30 min, followed by 30 min on ice. Phagocytic activity was analyzed in viable (DAPI-negative) cells by flow cytometry. Oxidative burst (in the presence of pHrodo™ 647 E. coli BioParticles) was assessed by measuring the fluorescence of dihydrorhodamine 123 (Merck).

For surface marker analysis, single-cell suspensions were stained with eBioscience Fixable Viability Dye (eBioscience, cat. no. 65-0865-14) and antibodies against CD63, CD66b, CD11b, CXCR2, TLR2, TLR4, and TLR5. Staining was performed for 30 min at 4°C, and fluorescence was analyzed by flow cytometry.

For intracellular cytokine staining, cells were treated with monensin (BioLegend, cat. no. 420701) and brefeldin A (BioLegend, cat. no. 420601) for 2 h. Cells were then stained with viability dye (eBioscience, cat. no. 65-0865-14), fixed, and permeabilized using the FoxP3 Fixation/Permeabilization Kit (BioLegend, cat. no. 421403) according to the manufacturer’s instructions. Subsequently, neutrophils and monocytes were stained with antibodies against IL-6, IL-8, IL-10, and IL-12. Fluorescence was measured by flow cytometry. Flow cytometric data were acquired using a BD Canto cytometer and analyzed with FlowJo software (v10).

## Results

### PN induces coordinated deployment of neutrophil-associated antimicrobial effectors

To define the molecular programs underlying renal innate immune activation in human PN, we compared FFPE tissue from PN nephrectomy specimens with control kidney tissue from healthy areas of renal cell carcinoma resections (Supplementary Table 1). HE-stained tissue sections indicated similar proportions of medullary and cortical tissue with more inflammation and reduced tubular integrity in PN (Figure 1A-B, Supplementary Figure 1). To identify molecular pathways driving antibacterial effector responses in PN, renal tissue was analysed by mass spectrometry–based proteomics, identifying over 2,500 proteins at FDR < 1% (Supplementary Table 2). Hierarchical clustering and principal component analysis (PCA) indicated clear separation of control and PN samples (Figure 1C, D). Gene Ontology Biological Process (GOBP) revealed coordinated activation of innate immune pathways consistent with antibacterial effector deployment. We observed upregulation of “*defense response to bacterium*” and “*antibacterial humoral response*”, which also includes “*AMP production*” (GO:0002775) (Figure 1E). At the same time, the proteins involved in mitochondrial electron transport and assembly of the mitochondrial respiratory chain were downregulated (Figure 1E). Cellular component analysis (GOCC) demonstrated preferential enrichment of extracellular and granule-associated compartments, including *“extracellular space”, “plasma membrane”, “azurophil granules”*, and *“Ig complex”* in PN (Figure 1F), consistent with neutrophil degranulation and AMP release. Further classification of the proteomic changes by REACTOME-based overrepresentation analysis showed upregulation of molecules involved in innate immunity, including factors of “*Neutrophil degranulation”*, “*Antimicrobial peptides”*, “*Extracellular matrix organization”* and proteins involved in “*Signaling by Interleukins”* (Figure 1G). Together, these data indicate a functional reprogramming of the renal proteome toward antimicrobial effector and degranulation pathways in human PN.

**Figure 1.**
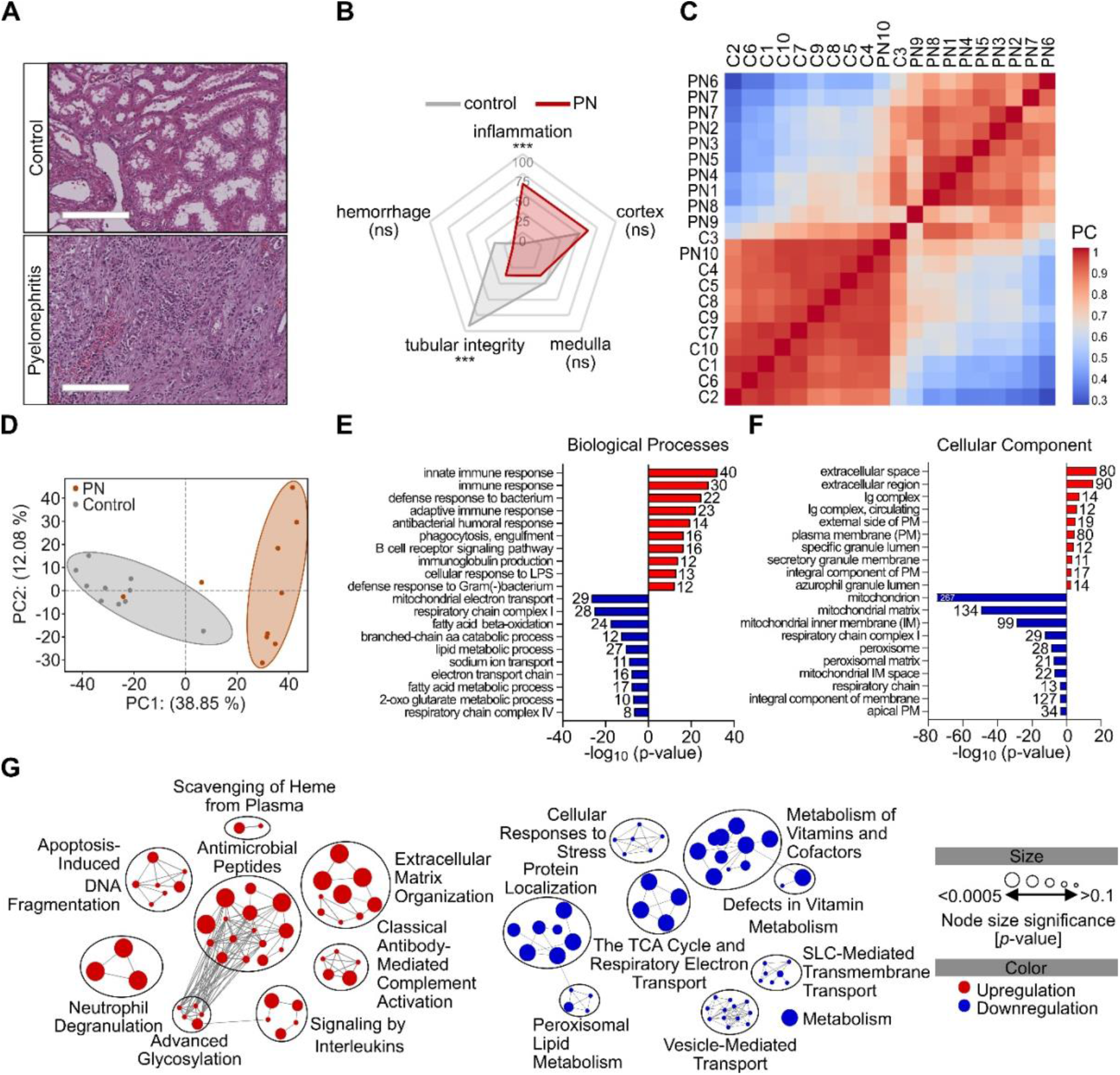
Proteomic alteration of the human kidney in PN. (**A, B**) Changes in inflammation, cortical and medullary tissue, tubular integrity, and haemorrhage were evaluated using a scoring sheet with an arbitrary scale of 0-100. The average scoring of PN (red) and control (grey) samples is visualized in the spider plot. \**p*<0.05, \*\*\**p*<0.0005. (**C-G**). Analysis of LC-MS/MS-based proteomics data of PN and control kidney sections. **(C)** Heatmap of Pearson correlation-based similarity between donors. The Pearson correlation cluster heatmap was generated using 1,273 proteins that were quantified in all 20 patients with an FDR-corrected *p*-value < 0.05. PC=Pearson correlation. (**D**) Principal component analysis (PCA) of the total kidney proteome of control and PN patients indicated condition-specific changes. (**E, F**) Enrichment analysis using Gene Ontology Biological Process (GOBP) (E) and Gene Ontology Cellular component (GOCC) (F). (**G**) REACTOME enrichment analysis by gene ontology of proteins expressed exclusively in one condition or with (log_2_(FC) > 2 or < −2, adjusted p-value < 0.05).

Given the enrichment of the REACTOME gene set “*Antimicrobial peptides*”, we next examined whether PN induces a coordinated increase in individual AMP effectors at the protein level. Of the 20 AMPs detected, 13 were significantly regulated (*p*-value < 0.05), including S100A12, S100A8, S100A9, CTSG, LCN2, LTF, RNASE3, BPI, AZU1, CAMP and other proteins involved in the host defence response (Figure 2A, B). Notably, resistin (RETN), recently described as an antimicrobial peptide^46^, was strongly induced in PN, whereas uromodulin (UMOD) was downregulated (Figure 2A, B). Thus, these data further expand the spectrum of antimicrobial effector molecules engaged during infection.

**Figure 2.**
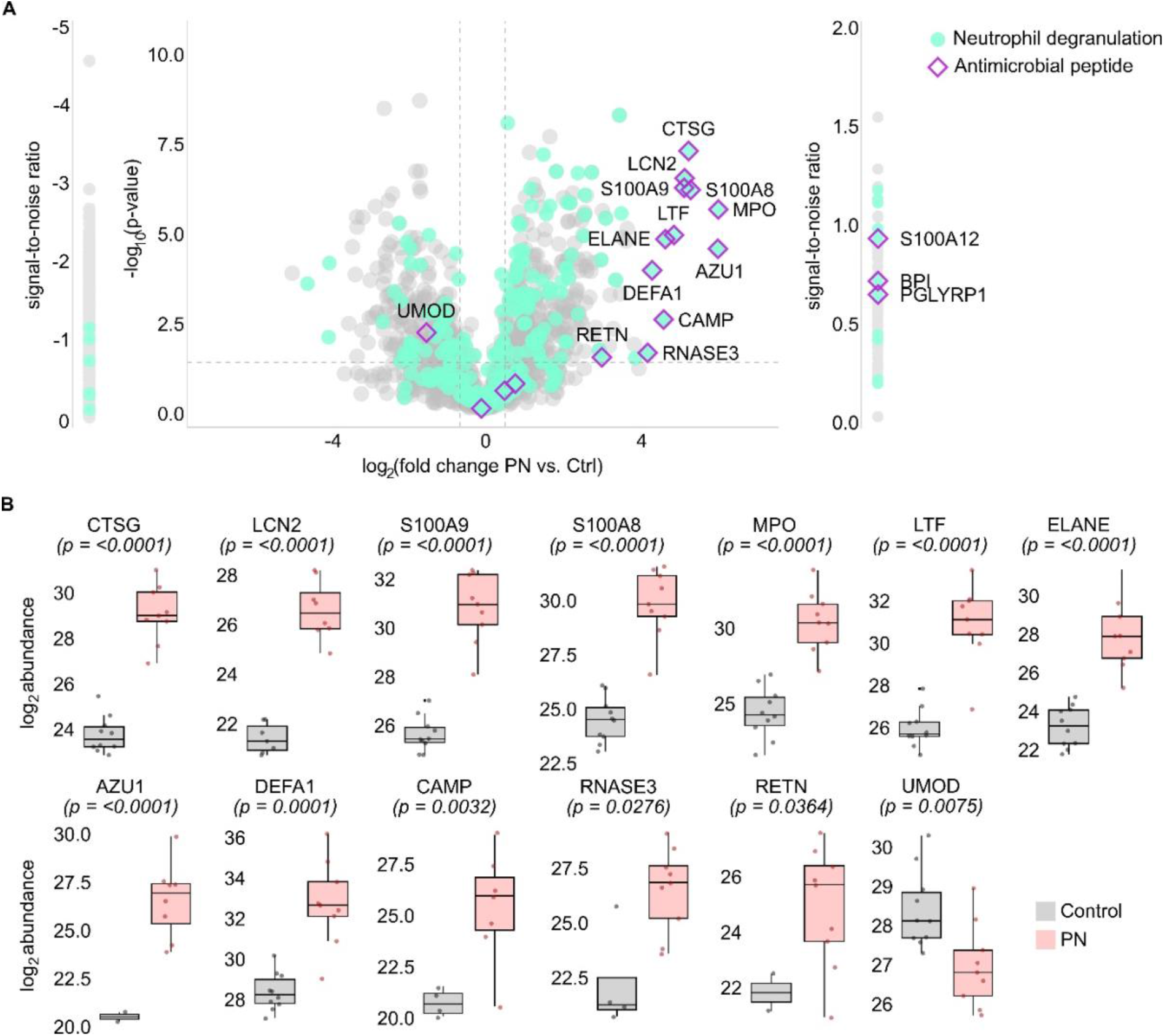
AMPs are more abundant in the PN kidney. (**A**) Volcano plot depicting the log_2_ fold change and −log_10_ *p-*value of proteins in PN versus control assessed by LC-MS/MS-based analysis of human kidney sections. Proteins detected only in one condition are indicated by the signal-to-noise ratio. (**B**) log_2_ protein abundance distribution of the 13 significantly (*p*<0.05) regulated AMPs detected in the kidney tissue of control (gray) and PN (red) patients.

### Urinary AMP upregulations recapitulate renal proteomic alterations

AMPs are primarily secreted into the renal tubular lumen and exert bactericidal activity at the site of infection in the renal pelvis and tubular system. Thus, urine proteomics provides a functional readout of renal AMP deployment into the tubular lumen, the site of inflammation during PN. Following this rationale, urine was collected from control donors (*n*=10) and PN patients (*n*=30), classified according to the ORENUC criteria for UTI stratification^28^ (Figure 3A). Age, sex, estimated glomerular filtration rate (eGFR), and leukocyte counts in urine and plasma were recorded (Table 1 and Supplementary Table 3).

**Figure 3.**
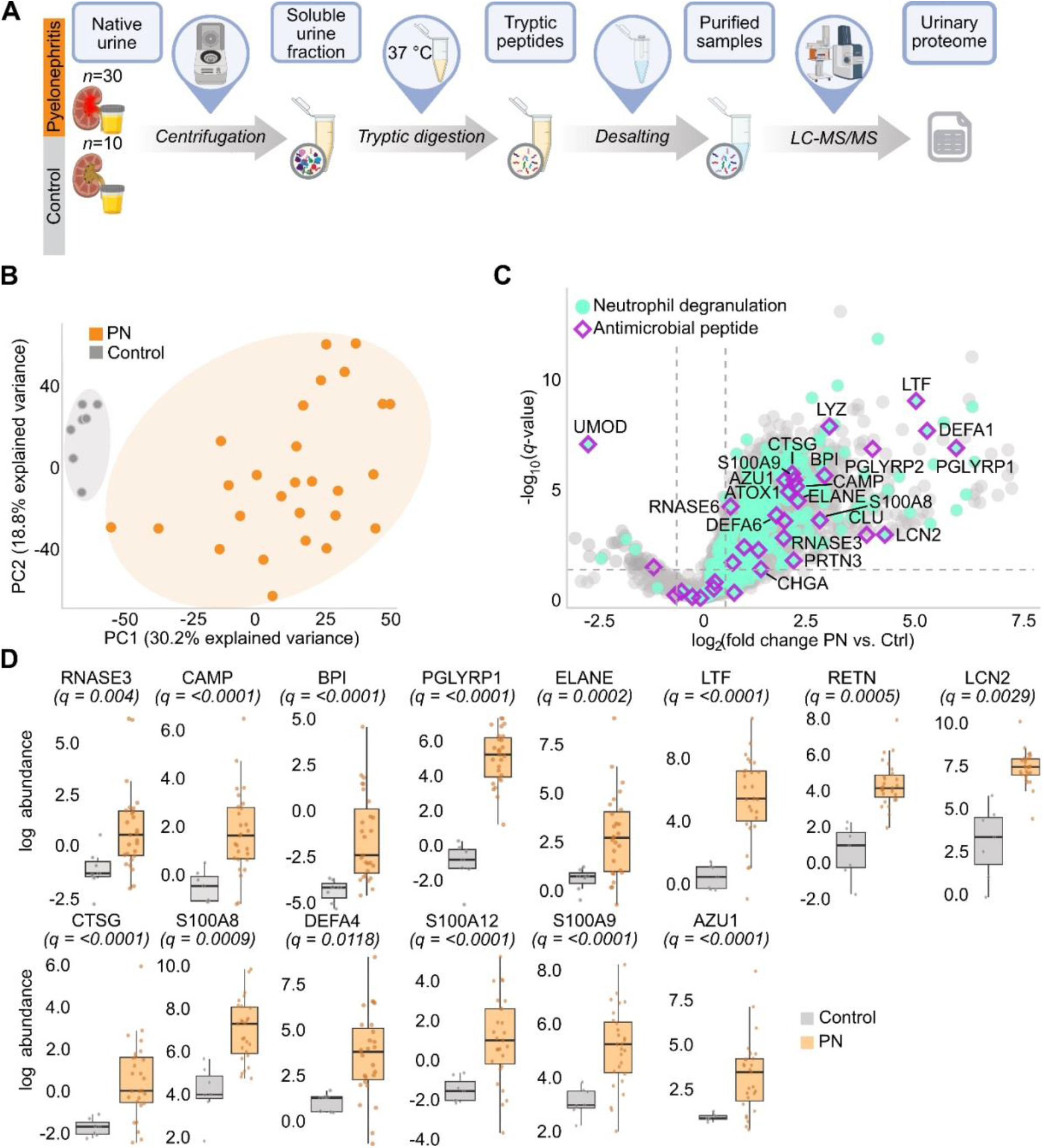
Significantly higher abundance of AMPs in the urine of PN patients. **(A)** Overview of the experimental pipeline. **(B)** Scatter plot of the first two principal components in principal component analysis (PCA), based on all proteins quantified (*n*=2,961) in the soluble urine fraction of PN patients and control donors (control). **(C)** Volcano plot showing log_2_ fold change difference between the soluble urinary proteome of PN patients versus controls. Proteins with a *q*-value (Student’s t-test, Benjamini-Hochberg FDR) <0.05 and a mean abundance difference >1.5 between PN and control groups were considered significantly regulated. Of the 1,716 proteins more abundant in PN patients, 23 of 33 associated with the REACTOME gene set “Antimicrobial peptides” are highlighted in magenta, and 243 of 343 proteins assigned to “Neutrophil degranulation” are shown in green. **(D)** Normalized log_2_ protein abundance distribution of antimicrobial proteins, defined as significantly regulated between the kidneys of PN and control patients in the soluble urine fraction.

**Table 1.** Characteristics and clinical data of the soluble urinary proteome of PN and control. (Related to Supplementary Table 3) eGFR = estimated glomerular filtration rate, CRP-C-reactive protein, nd = not determined.

| Parameters | PN | Control |
| --- | --- | --- |
| N | 30 | 10 |
| Mean age (min, max) | 38 (16 – 86) | 29 (23 – 31) |
| Sex (n) | ♀: 26; ♂: 4 | ♀: 6; ♂: 4 |
| eGFR | 101.6 (27.8 – 168.2) | nd |
| Fever [°C] | 39 (37.8 – 41.2) | nd |
| CRP [mg/mL] | 102.7 (13.7-280.2) | nd |
| Leukocytosis [cells/mL plasma] | 13.8 (8.9 – 25) | nd |
| Leukocyturia [cells/mL urine] | 2289.6 (31 – 33748) | nd |

Using LC-MS/MS, we quantified 2,961 proteins in the urine proteome, with differential expression in PN and control samples, indicated by segregation of the samples using dimensionality reduction (Figure 3B and Supplementary Table 4). Comparative analysis defined 1,743 significantly altered proteins (*q*-value, Benjamini-Hochberg < 0.05; fold change > 1.5), including strong enrichment of proteins associated with the gene sets “*Antimicrobial peptides*” and “*Neutrophil degranulation*” in urine of PN patients (Figure 3C). Notably, proteins previously described in kidneys of late-stage PN patients were also upregulated, suggesting their release into the tubular lumen (Figure 3D). AMP abundances positively correlated with urine leukocyte counts, with hierarchical clustering revealing coordinated regulation of S100A12, LTF, BPI, ELANE, cathepsin G (CTSG), and the S100A8/A9/DEFA4 cluster. Upregulation of selected AMPs and UMOD was further validated by ELISA (Supplementary Figure 2), supporting a direct link between immune cell recruitment and shaping of the antimicrobial composition of the urinary microenvironment.

### AMPs colocalize with myeloid cells in kidney tissue

In our study, we observed strong clustering of leukocytes with AMPs (Supplementary Figure 2A), indicating that leukocytes are primary source of AMPs during PN. To analyse AMP and immune cell distribution, we performed multiplex immunofluorescence microscopy using sequential staining of S100A8, azurocidin-1 (AZU1), CTSG, and S100A12; immune cell markers CD14 and CD66b; and the tubular markers AQP1 and NKCC2. Following cell segmentation, CD66b^+^cells were annotated as neutrophils, and CD66b^−^ CD14^+^cells as monocytes and monocyte-derived macrophages. S100A8, AZU1, CTSG, and S100A12 showed strong spatial colocalization with neutrophils, monocytes and macrophages (Figure 4A, B). Notably, AQP1^+^ and NKCC2^+^ tubules showed no specific spatial colocalization with AMPs, identifying myeloid cells as dominant sources of AMP effectors during PN.

**Figure 4:**
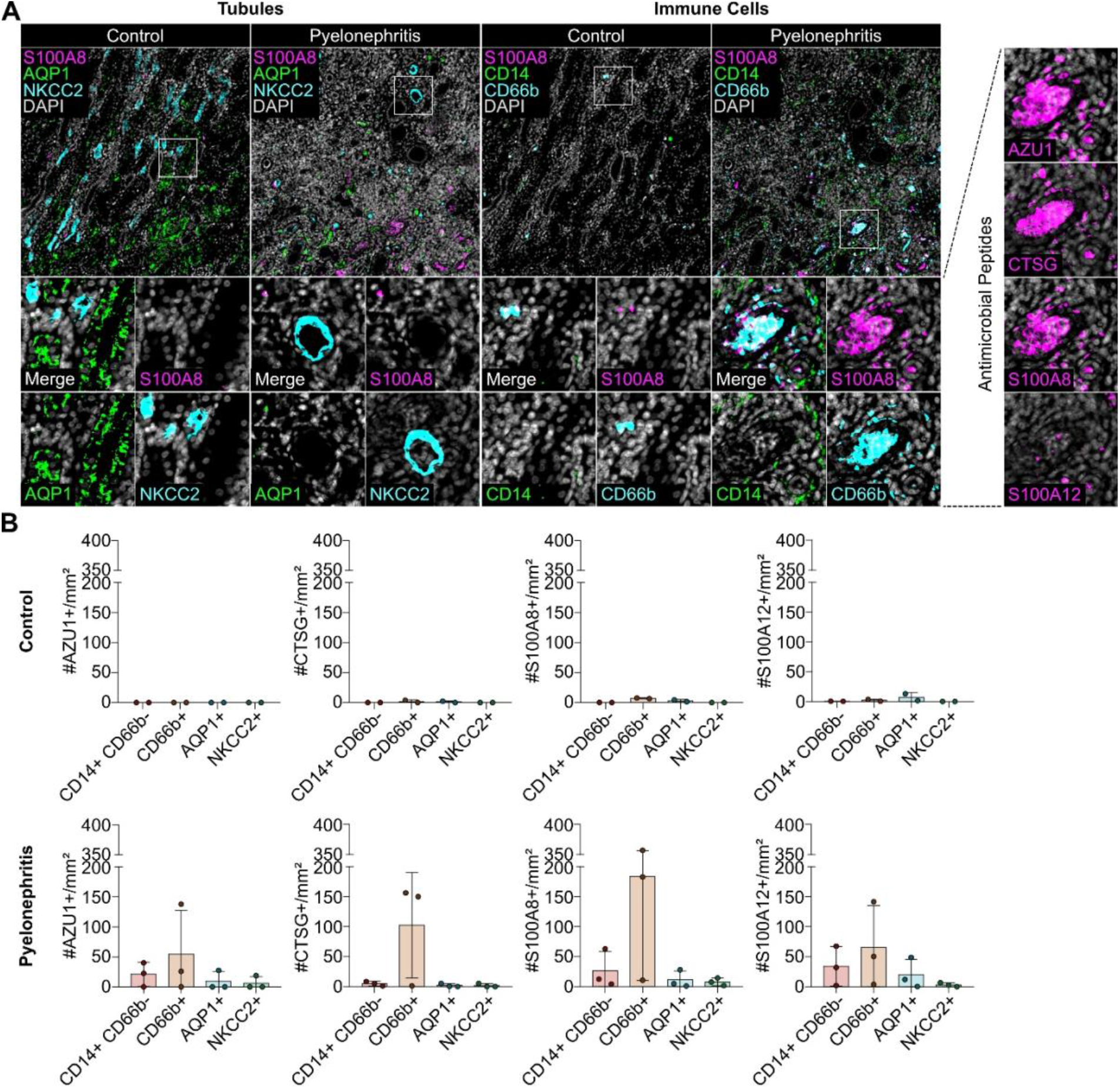
Myeloid cells represent the primary source of AMPs in human PN. FFPE human kidney sections from PN (*n*=3) and control patients (*n*=2) were stained with DAPI (nuclei), CD14 (monocytes), CD66b (neutrophils), AQP1 (proximal tubules), NKCC2 (distal tubules), and the following AMPs: AZU1, CTSG, S100A8, and S100A12. **(A)** Exemplary images of immune and tubular markers as well as AMPs in kidney tissue from patients C2 (Control) and PN3 (Pyelonephritis) **(B)** Frequency of AMP^+^ CD14^+^ CD66b^-^ monocytes (red), AMP^+^ CD66b^+^ neutrophils (orange), AMP^+^ AQP1^+^ stromal cells (turquoise), and AMP^+^ NKCC2^+^ stromal cells (green) per mm^2^ in control (top) and PN (bottom) kidney tissue.

### Urinary peptidomics uncovers novel EPs in PN patients

Beyond classical AMPs, proteolytic processing of immune-derived precursor proteins can generate EPs, adding a post-translational layer to antimicrobial regulation^9^. To identify novel EPs in the urine of PN patients, we performed native peptidomics, enabling unbiased detection of proteolytically generated peptide sequences (Figure. 5A). In total, 9,295 peptides were identified at a stringent FDR (<0.01), of which 8,504 were unique to the PN cohort (Supplementary Table 5). Application of six AMP prediction algorithms^36–41^ indicated 10 sequences with putative antimicrobial function, each supported by at least five out of six classifiers (Fig. 5B). None of these candidates have previously been linked to PN. Among them, one sequence corresponded to a known antimicrobial peptide, Calcitermin (VAIALKAAHYHTHKE), derived from the C-terminus of S100A12. As S100A12 was elevated in the kidney tissue and urine of PN patients, we examined Calcitermin expression in detail and confirmed peptide identity by robust b- and y-ion series using LC-MS/MS (Fig. 5C). PRM-based LC-MS quantified Calcitermin at 20.3 fmol/µg and 53.2 fmol/µg native peptides in urine from two PN patients (Figure. 5D and Supplementary Table 6A, B), confirming its presence *in vivo*. Given previous reports of Calcitermin’s antimicrobial activity under acidic conditions, we tested its effect against UPEC. Calcitermin markedly inhibited bacterial growth only at acidic pH 5.0 (Fig. 5E), but not at neutral pH 7.0 (Supplementary Figure 3). Since AMPs, including EPs, function not only as direct microbicidal agents but also as modulators of leukocyte responses, we tested their protective capacity beyond pathogen killing.

**Figure 5.**
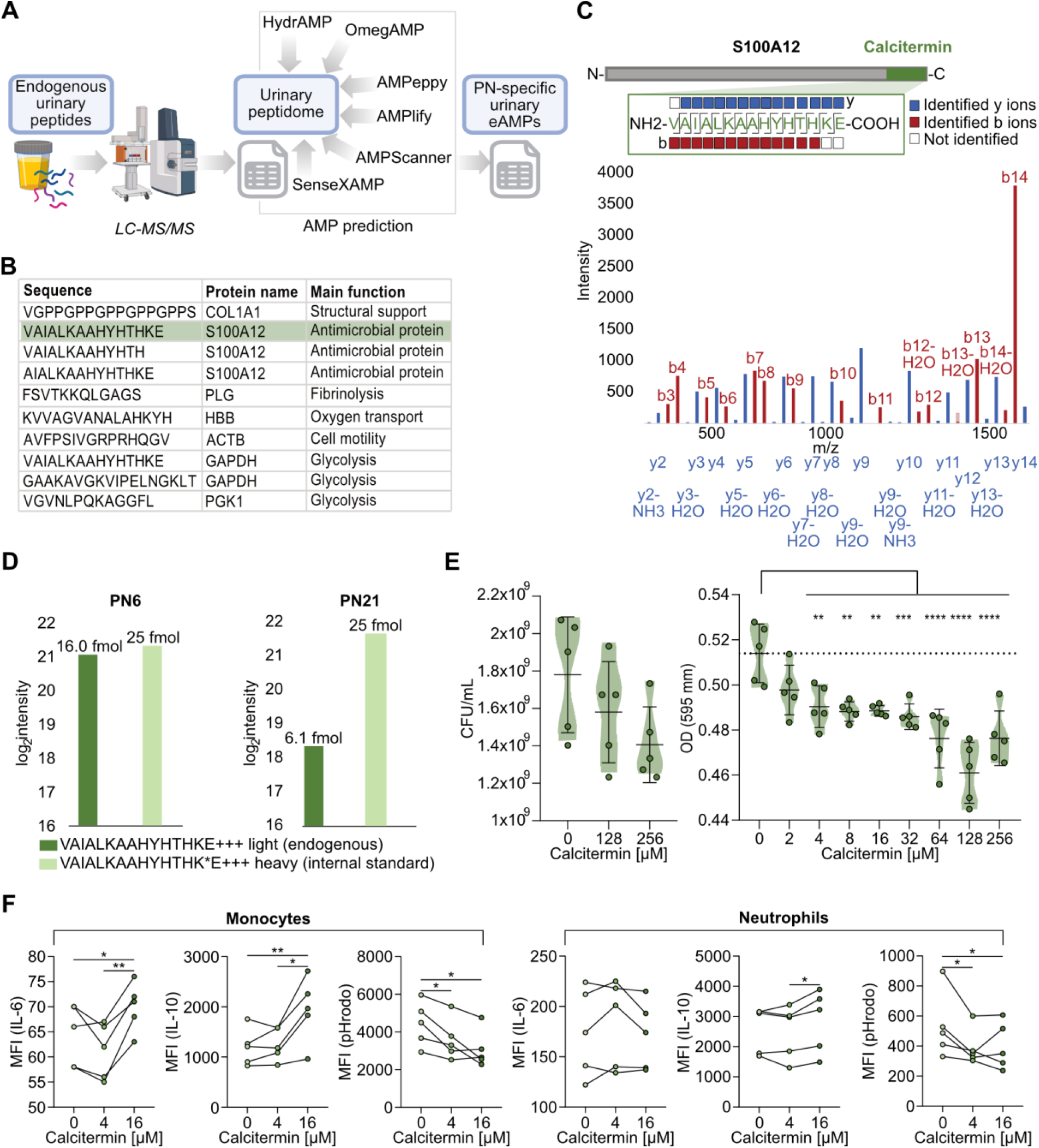
Urine peptidomics revealed encrypted antimicrobial peptides (EPs) with immunomodulatory effects. (**A)** Overview of eAMP prediction from the native urinary peptidome of PN patients. (**B)** PN-specific endogenous peptides with a classifier score >5 in the LC-MS/MS dataset, predicted to be antimicrobial by at least 5 out of 6 AMP predictors. (**C)** Identified b and y ions of endogenous Calcitermin in LC-MS/MS data of patient PN6 (563.645 m/z; Charge 3). **(D)** Absolute quantification of endogenous Calcitermin and heavy-labeled synthetic Calcitermin (internal standard) in the urine of selected PN patients by parallel reaction monitoring (PRM)-based LC-MS/MS. For PN6, the intensity corresponds to 53.3 fmol/ug native peptides, for PN21 to 20.3 fmol/ug native peptides. **(E)** Colony-forming units per millilitre (CFU/mL) and optical density (OD) at 595 nm of UPEC 536, supplemented with titrated concentrations of Calcitermin for 24 hours at pH 5.0. **(F)** Neutrophils and CD14+ monocytes were isolated from the peripheral blood of healthy human donors (*n*=5) and exposed to 0, 4, and 16 μM Calcitermin for 18 h. Dot plots for monocytes and neutrophils depict the mean fluorescence intensity (MFI) of fluorophore-conjugated antibodies targeting interleukin-6 (IL-6) and IL-10 as well as pHrodo Bioparticles conjugates. \**p*<0.05, \*\**p*<0.005, \*\*\**p*<0.005, \*\*\*\**p*<0.0005.

To investigate the immunomodulatory potential of Calcitermin, human leukocytes isolated from healthy donors were incubated with the peptide at 4 µM (the minimal concentration eliciting a significant antibacterial effect) and 16 µM (a prospective application-relevant concentration) for 18 h. Calcitermin showed no detectable cytotoxicity toward primary human cells (Supplementary Fig. 4), while selectively inducing IL-6 and IL-10 production in monocytes, but not in neutrophils (Fig. 5F). Notably, Calcitermin also induced a concentration-dependent reduction in phagocytic activity in both monocytes and neutrophils (Fig. 5F).

Together, these findings demonstrate that PN-associated proteomic remodelling generates endogenous and functionally active EPs, exemplified by Calcitermin, which retains antibacterial activity against UPEC under infection-relevant conditions and modulates myeloid cell function, linking antimicrobial activity to regulation of leukocyte function and suggesting a dual role in shaping host defence during infection.

## Discussion

Pyelonephritis induces a compartmentalized antimicrobial defence program in which immune cell–derived precursor proteins are spatially deployed and proteolytically processed into bioactive peptides within the renal parenchyma and urinary microenvironment. Our data support a model in which leukocyte recruitment, granule release, and local proteolysis together shape the antimicrobial landscape at the site of infection.

At the tissue level, we observed extensive remodelling of the renal proteome in PN, characterized by coordinated activation of innate immune pathways, neutrophil degranulation, and extracellular matrix organization. These findings are consistent with experimental models demonstrating that myeloid cell recruitment drives bacterial containment, while simultaneously promoting tissue inflammation and remodeling^47^. Within this context, we defined robust upregulation of classical AMPs, including S100A12, S100A8/S100A9, CTSG, AZU1, BPI, LCN2, and LTF, which together constitute a core neutrophil antimicrobial module. These molecules act through complementary mechanisms, including metal sequestration, membrane disruption, and enzymatic degradation, to restrict bacterial growth and modulate inflammatory signalling. The concerted induction of this AMP repertoire highlights neutrophil effector deployment as a central mechanistic axis of host defence in human PN.

In parallel, we observed downregulation of UMOD, a highly abundant tubular protein with established roles in modulating neutrophil activation, complement binding, and bacterial aggregation^48,49^. Reduced UMOD abundance likely reflects tubular stress or impaired secretion during infection and may functionally shift the balance toward heightened immune activation within the tubular lumen. Conversely, the strong induction of RETN, recently recognized as a proinflammatory and bactericidal AMP^46,50^, further expands the spectrum of antimicrobial effectors secreted during PN. Together, these findings indicate that PN reshapes the urinary proteome through both immune cell–derived effector functions and altered epithelial contributions, collectively tuning the antimicrobial and inflammatory milieu of the urinary tract.

Spatially resolved multiplex immunofluorescence microscopy provided novel insights into the cellular origin of the AMP response. We defined strong spatial colocalization of S100A8, S100A12, CTSG, and AZU1 with CD66b^+^ neutrophils and CD14^+^monocytes/monocyte-derived macrophages, establishing myeloid cells as dominant sources of AMP effector proteins during infection. While neutrophils are well-established producers of S100 proteins, cathepsins, and defensins, the observed contribution of monocytes and macrophages underscores a broader, multicellular antimicrobial network. In parallel, epithelial cells retain the capacity to express CAMP^51^, LTF^52^, and LCN2^20^, while monocytes and macrophages are capable of producing S100A8 and S100A9^53,54^. These data suggest that epithelial and immune compartments cooperate to establish layered antimicrobial protection. Thus, the AMP response in PN emerges as a coordinated defence program integrating leukocyte recruitment, cellular specialization, and spatial compartmentalization.

A key novelty of our study is the demonstration that this AMP-rich environment supports the *in vivo* generation of EPs. Native urinary peptidomics suggest extensive proteolytic processing of host proteins during infection, uncovering multiple peptide fragments with predicted antimicrobial activity. Machine learning–based AMP prediction^36–41^ identified ten of these proteolytic peptides to exert antimicrobial activity^9,55^. Among these, the S100A12-derived peptide Calcitermin was particularly compelling, as it was detected and quantitatively confirmed in PN patient urine and exhibited antibacterial activity against UPEC under acidic conditions that mirror the inflamed urinary tract. Structurally, Calcitermin is enriched in histidine and lysine residues, conferring cationic charge and zinc-binding capacity that promote membrane interaction and bacterial killing in low-pH environments. The concomitant upregulation of its precursor protein, S100A12, in both renal tissue and urine supports a model whereby infection-induced proteolytic activity converts abundant immune-derived proteins into functionally active antimicrobial fragments, consistent with the encrypted immunity hypothesis^55^.

EPs are increasingly recognized as multifunctional effectors that integrate direct microbicidal activity with the regulation of host immune responses. Previous studies on fibrinogen- and apolipoprotein-derived EPs have demonstrated that such peptides can shape leukocyte activation, cytokine production, and inflammatory resolution in both *in vitro* and i*n vivo* infection models ^56 57–59^. In this context, our data identify Calcitermin as a peptide with a comparable dual mode of action, combining antibacterial efficacy with selective immunomodulatory effects on primary human immune cells. The induction of IL-6 and IL-10 in monocytes suggests a coordinated activation of pro- and anti-inflammatory pathways, potentially enabling a balanced immune response. At the same time, the observed reduction in phagocytic activity, in both monocytes and neutrophils, may reflect a regulatory mechanism that limits excessive cellular activation and tissue damage.

In PN, such combined antimicrobial and immunomodulatory properties could have system-level consequences by simultaneously reducing bacterial burden and modulating inflammatory responses within the kidney. This dual activity could limit the transition from acute infection to chronic inflammation, thereby preserving tissue integrity and function. Moreover, modulation of leukocyte activity could influence the recruitment, activation state, and lifespan of innate immune cells within infected tissue. On a broader level, peptides like Calcitermin may contribute to restoring immune homeostasis by fine-tuning the balance between pathogen clearance and host-mediated damage. Together, these findings support the concept that EPs act not merely as antibiotics but as regulators of host defence, with potential therapeutic relevance in complex infections such as pyelonephritis. In the context of rising multidrug resistance among uropathogens, peptides such as Calcitermin may provide a blueprint for therapeutic strategies that harness or mimic innate defence mechanisms in addition to directly targeting bacterial viability. Moreover, delineating the proteases and microenvironment conditions that govern EP generation may uncover novel pharmacological targets to enhance peptide production, stability and activity *in vivo*.

In summary, this study provides an integrative, mechanistic framework for understanding AMP/EPs-mediated host defence in human PN. By linking immune cell recruitment to AMP deployment, proteolytic processing, and context-dependent antibacterial activity, we define a multi-layered antimicrobial defence program operating across renal tissue and urine. These findings expand current concepts of innate immunity in the urinary tract and establish a foundation for future studies aimed at identifying the proteases responsible for EPs generation, validating AMP/EPs signatures in larger cohorts, and exploring peptide-based interventions for urinary tract infections.

## Supporting information

Supplementary Table 1 - Clinical Data of Control and Pyelonephritis Patients

Supplementary Table 2 - LC-MSMS-based Proteomics of Kidney Pyelonephritis

Supplementary Table 3 - Clinical Data of Control Urine Donors and Pyelonephritis Patients

Supplementary Table 4 - LC-MSMS-based Urine Proteomics of Pyelonephritis

Supplementary Table 5 - AMP Classification for Identified Urinary Native Peptides

Supplementary Table 6 - Parallel Reaction Monitroring (PRM) Mode LC-MSMS-based Absolute Quantification

## Data availability

The raw mass spectrometry proteomics data have been deposited to the ProteomeXchange Consortium http://proteomecentral.proteomexchange.org via the PRIDE partner repository^60^ with the dataset identifiers PXD042016 for kidney proteome data, PXD069781 for urinary proteome data and PXD069829 for urinary peptidome data.

The supplementary tables are accessible via the following link: https://uni-duisburg-essen.sciebo.de/s/MLQsjW377b9PkyL

## Acknowledgements

We acknowledge support from the Open Access Publication Fund of the University of Duisburg–Essen. We received funding from the German Research Foundation: FOR5427/1- (466687329): FOR5427 SP1 (OS; FW); FOR5427 SP3 (UD) FOR5427 SP4 (DRE); FOR5427 SP7 (SvV); : 247377969 (HS), EN984/16-1, 18-1, 19-1 (DRE); and TR332 A3 and Z1 (DRE), INST 20876/486-1 (DRE), and from the European Union: ERA-NET NEURON (01EW2503) (OS). Furthermore, we thank Edda Christians from Hannover Medical School and Kerstin Wilhelm and Tanja Bloch from Justus-Liebig University Gießen for their technical assistance and valuable contributions to the study. We also thank the Imaging Center Essen (IMCES) at the Faculty of Medicine, University of Duisburg–Essen, Germany, for providing access to the AxioScan.Z1, instrument maintenance, and general support with immunofluorescence microscopy.

---

**Supplementary Figure 1:**
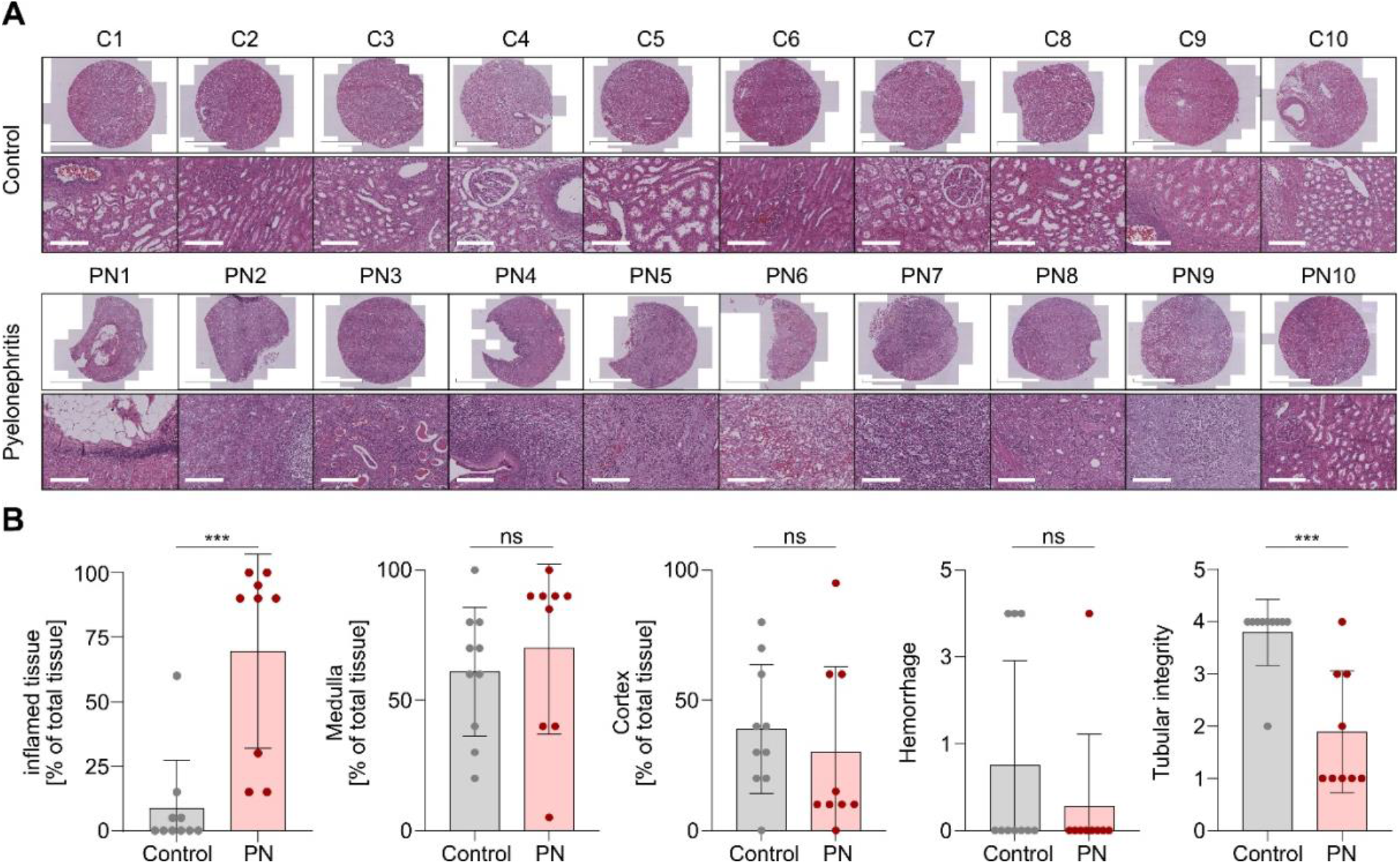
Evaluation of PN and control kidney tissue integrity. **(A)** Images of H&E-stained kidney tissue from control and PN patients. **(B)** Evaluation of tissue stainings according to the proportion of inflamed, medullary, and cortical tissue, hemorrhage, and tubular integrity. Related to Figure 1A, B. *=*p*<0.05, \*\*\**p*<0.0001, *ns*=non-significant.

**Supplementary Figure 2.**
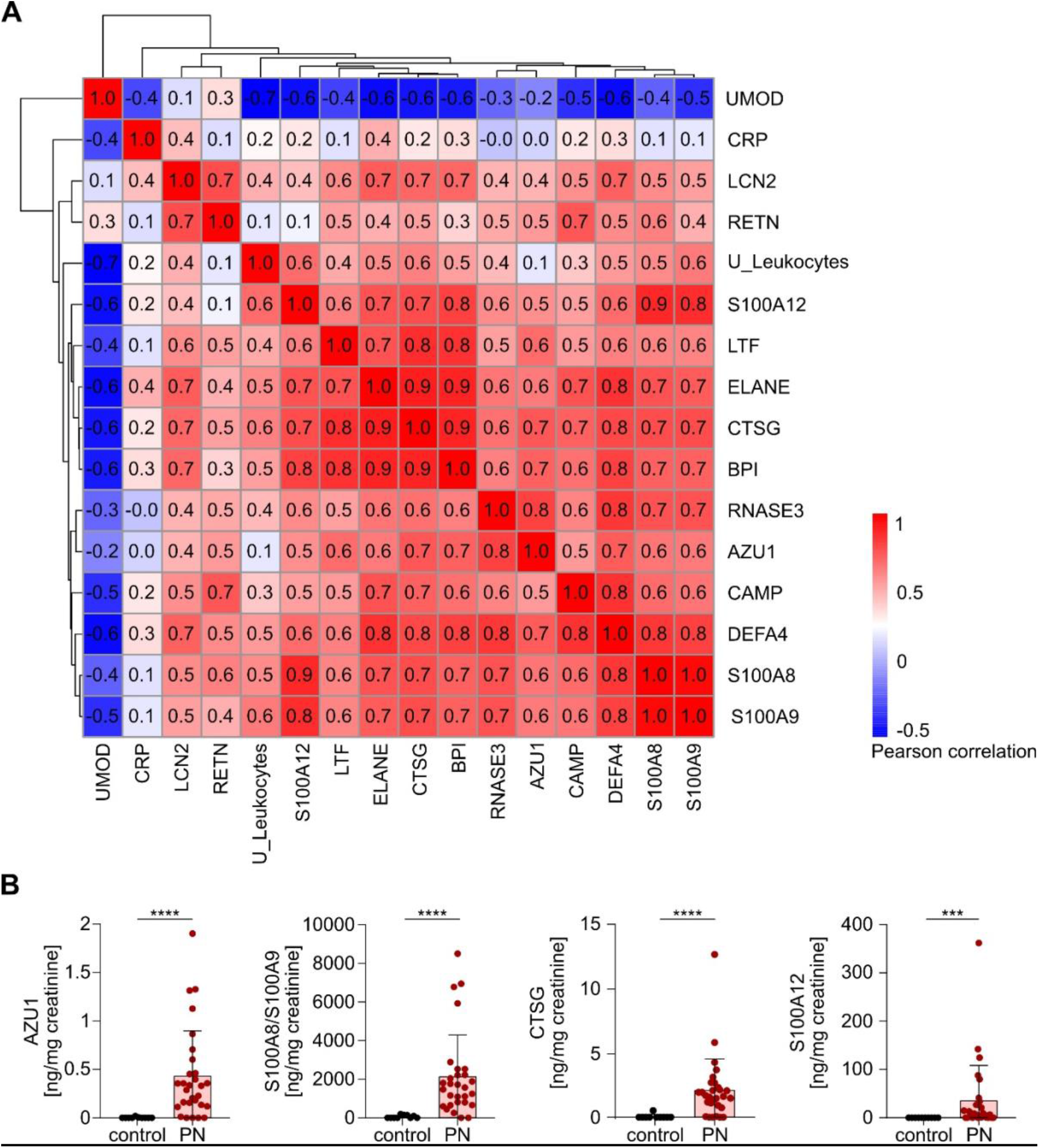
Regulation of AMPs in the urine of acute PN patients. (**A**) Hierarchical clustering using Ward.D linkage, based on Pearson correlation coefficients calculated between the normalized log_2_-transformed protein abundances of selected AMPs (as determined by LC–MS in urine samples from patients with acute PN and control healthy donors) and the urinary leukocyte count. **(B)** Scatter plots depicting the concentration of AZU1, S100A8/S100A9, CTSG, and S100A12 in urine from patients with acute PN (*n*=29) and controls (*n*=10), measured by ELISA, and normalized to urine creatinine \*\*\**p*<0.005, \*\*\*\**p*<0.0001.

**Supplementary Figure 3.**
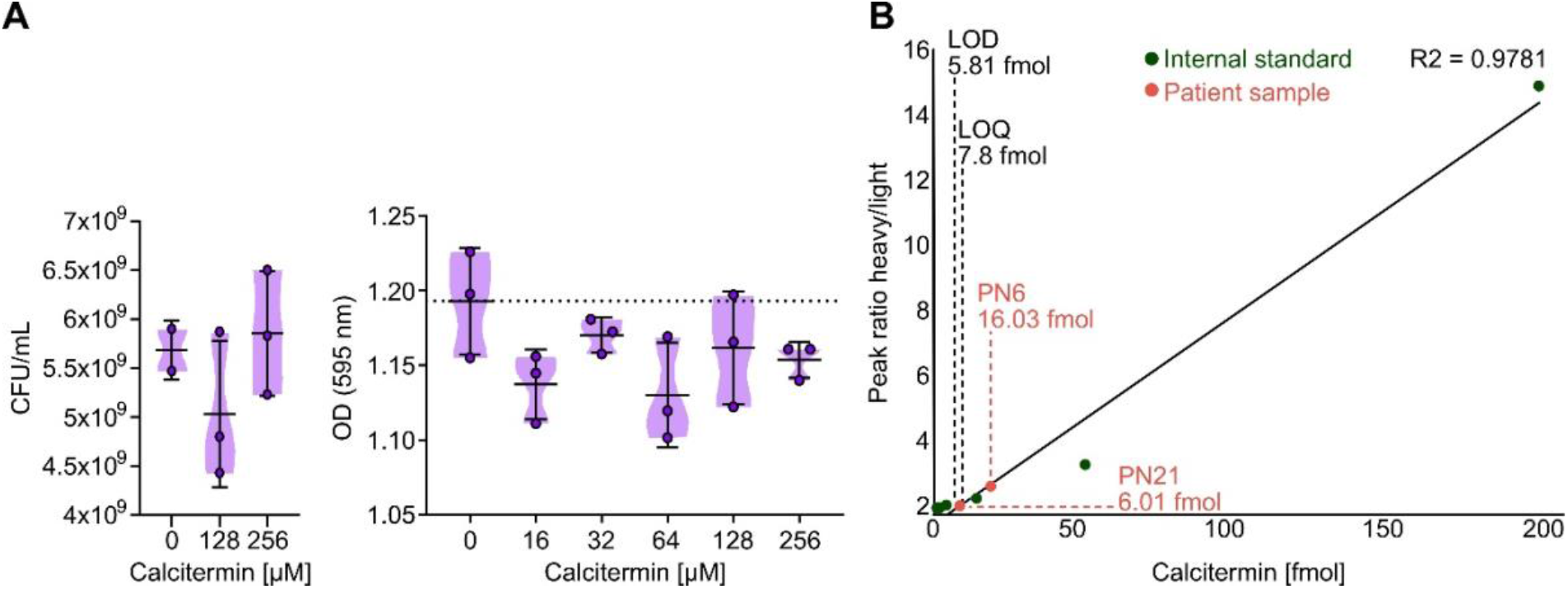
Antimicrobial activity and quantitative detection of Calcitermin in urine. (**A**) Colony-forming units per millilitre (CFU/mL) and optical density (OD) at 595 nm of UPEC 536, supplemented with titrated concentrations of Calcitermin for 24 hours at pH 7.0 **(B)** Calibration curve for Calcitermin, covering a range of 3.125–200 fmol. The determined concentration of endogenous Calcitermin in patient samples with a Calcitermin concentration > LOD is indicated.

**Supplementary Figure 4.**
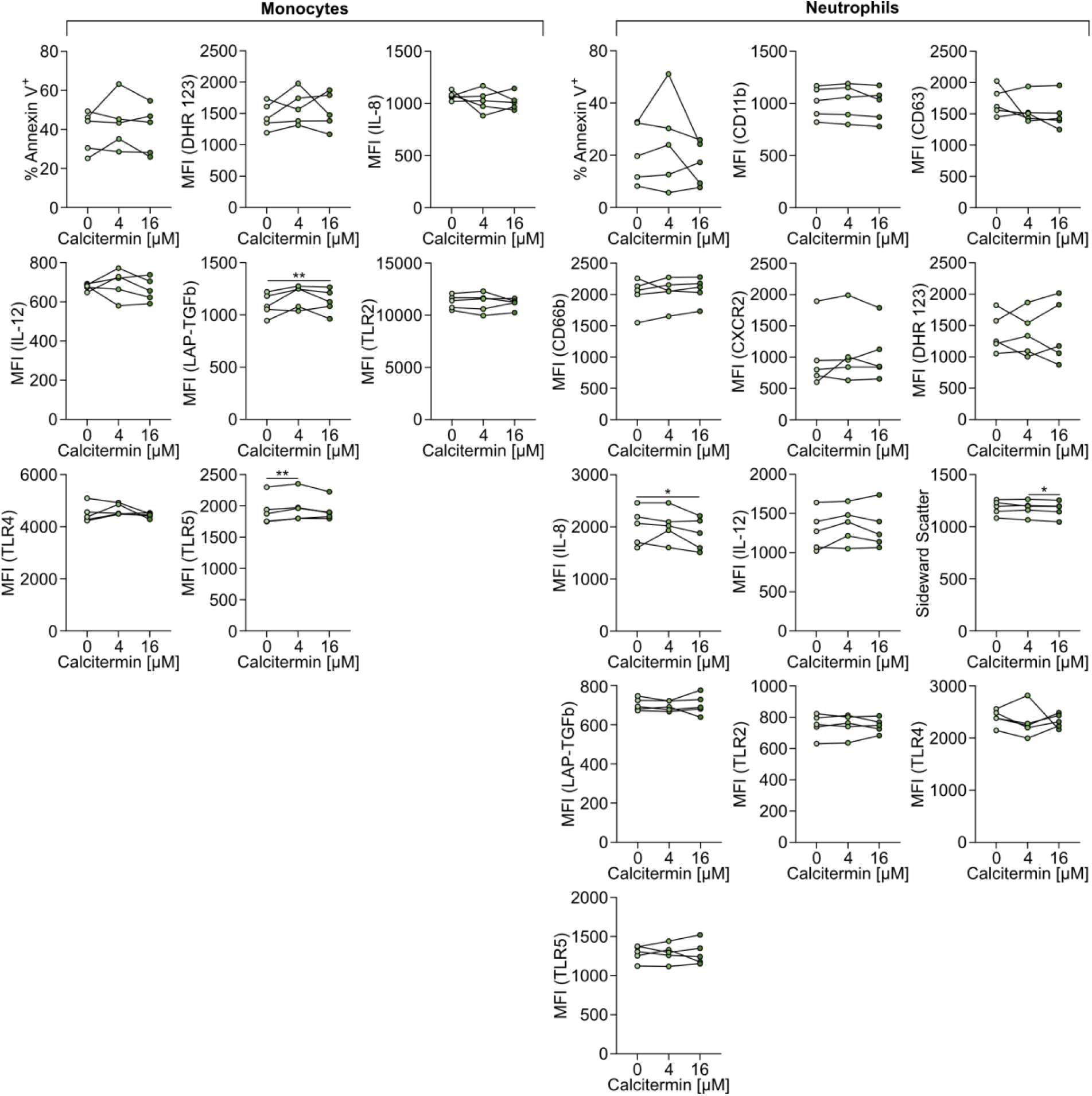
Flow cytometric analysis of monocytes and neutrophils following Calcitermin treatment. Neutrophils and CD14+ monocytes were isolated from peripheral blood of healthy donors (n = 5) and incubated with 0, 4, or 16 μM Calcitermin for 18 h. Representative dot plots show side scatter characteristics, the proportion of Annexin V+ apoptotic cells, and mean fluorescence intensity (MFI) of markers associated with activation and function, including IL-8, IL-12, CD11b, CD63, CD66b, CXCR4, LAP–TGF-β1, TLR2, and TLR4. Reactive oxygen species (ROS) production was assessed using dihydrorhodamine 123 (DHR 123). Statistical significance is indicated as *p < 0.05 and **p < 0.005.

